## Supplementary Information for "IM30 IDPs form a membrane protective carpet upon super-complex disassembly"

***Address correspondence to**

### Supplementary Data and Figures

*IM30 binding to membranes involves structural rearrangement*

We analyzed possible structural alterations upon membrane binding of IM30 via tryptic digestion of IM30 in absence and presence of DOPG liposomes. The protein (2.5 µM with or without liposomes (0.1 mM DOPG)) was incubated for 30 min at 4 °C, and then the protease was added (0.01 mg/mL) (Supplementary Fig. 1). Samples were taken at the indicated time points and analyzed via SDS-PAGE. As shown previously, helix 7 of IM30 is rapidly cleaved[1,2]. The remaining fragment H1-6, *i.e.* the PspA-domain, is more protease-resistant, and thus the corresponding band is detectable until about 10 min after addition of trypsin. Strikingly, in contrast to soluble IM30, this band remained visible for about 60 min after trypsin addition when DOPG liposomes were present (Supplementary Fig. 1, red box). Thus, membrane binding appears to protect the PspA-domain from tryptic digestion. The altered protease resistance in presence of DOPG membranes indicates a structural change of IM30 upon membrane binding, which involves shielding of protein regions.

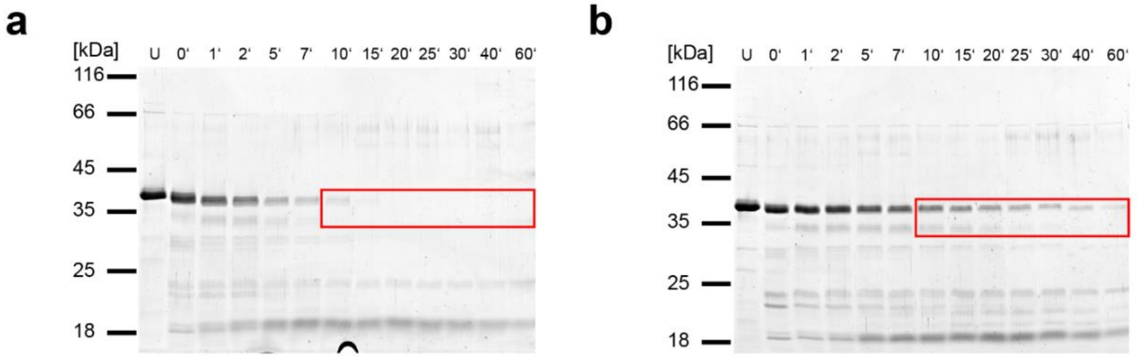


***Supplementary Figure** 1*: **The structure of IM30 changes upon membrane binding.**

**(a/b)** Limited trypsin digestion of IM30 in absence (a) and presence (b) of DOPG liposomes shows that the protease accessibility of IM30 is altered upon binding to PG liposomes. U=untreated sample. Time points indicate the time of incubating the sample with trypsin. The boxed area highlights the bands showing the major difference between trypsin digestion in absence *vs*. presence of DOPG.

*A homo-dimeric IM30**

IM30 spontaneously forms diverse ring structures with different rotational symmetries[3–8]. Furthermore, individual rings appear to stack and form rod-shaped structures[1,3–6,9,10]. Because of this pronounced intrinsic structural heterogeneity, which complicates structural and functional analyses, it was necessary to homogenize the sample via destabilizing the compact structure of IM30 rings. As described previously, this can *e.g.* be achieved via mutation of a highly conserved amino acid cluster in helix 4 (see Supplementary Fig. 2a)[3]. The resulting mutant, IM30_FERM, readily forms tetramers instead of rings[3]. However, we noticed an increased tendency of this mutant to unspecifically aggregate[3], and thus this variant turned out to be unsuitable for further structural characterization. Hence, we aimed to create a more stable version of this mutant by mutational surface engineering. Following the rationale described by Derewenda[11], we reduced the local conformational entropy by replacing large hydrophilic residues at the protein surface with Ala. We selected Glu83 and Glu84 as promising candidates for mutation, which are located in a predicted loop region (Supplementary Fig. 2a), as loop regions with a high abundance of acidic/basic residues are likely solvent-exposed and thus located at the protein surface[12]. Indeed, the resulting mutant, IM30* (IM30_FERM_EE), was less prone to aggregation, even after prolonged incubation in HEPES buffer (Supplementary Fig. 2b). Additionally, we noticed that increasing the ionic strength of the buffer by the addition of NaCl strongly affected the oligomeric state of IM30* (Supplementary Fig. 2b). A small increase in the ionic strength by the addition of 50 mM NaCl leads to nearly complete clustering of the protein and formation of aggregates with molecular masses > 300 kDa. Further addition of NaCl resulted in formation of dimers (~60 kDa) that were stable at NaCl concentrations of at least up to 300 mM. This shift in the oligomeric state did also affect the thermal stability of the protein, as measured via CD-spectroscopy (Supplementary Fig. 2c). Increasing the NaCl concentration from 0 to 300 mM resulted in an overall shift of the melting temperature $T_m$ from 53.2±0.7 °C to 48.0±0.1 °C. Importantly, addition of NaCl did not affect the secondary structure of the protein (see Supplementary Fig. 1e/f). Thus, the changes in $T_m$ was most likely exclusively caused by the change in the oligomeric state, accounting for the overall compactness of the structure or the surface accessibility, respectively, which is a major constituent for protein stability against denaturation[13,14]. As depicted in Supplementary Fig. 1d, the observed changes in $T_m$ nicely correlate to the change in the oligomeric state. In summary, IM30* appeared to be perfectly suitable for further structural and functional analyses, as it exclusively forms stable dimers at defined conditions.

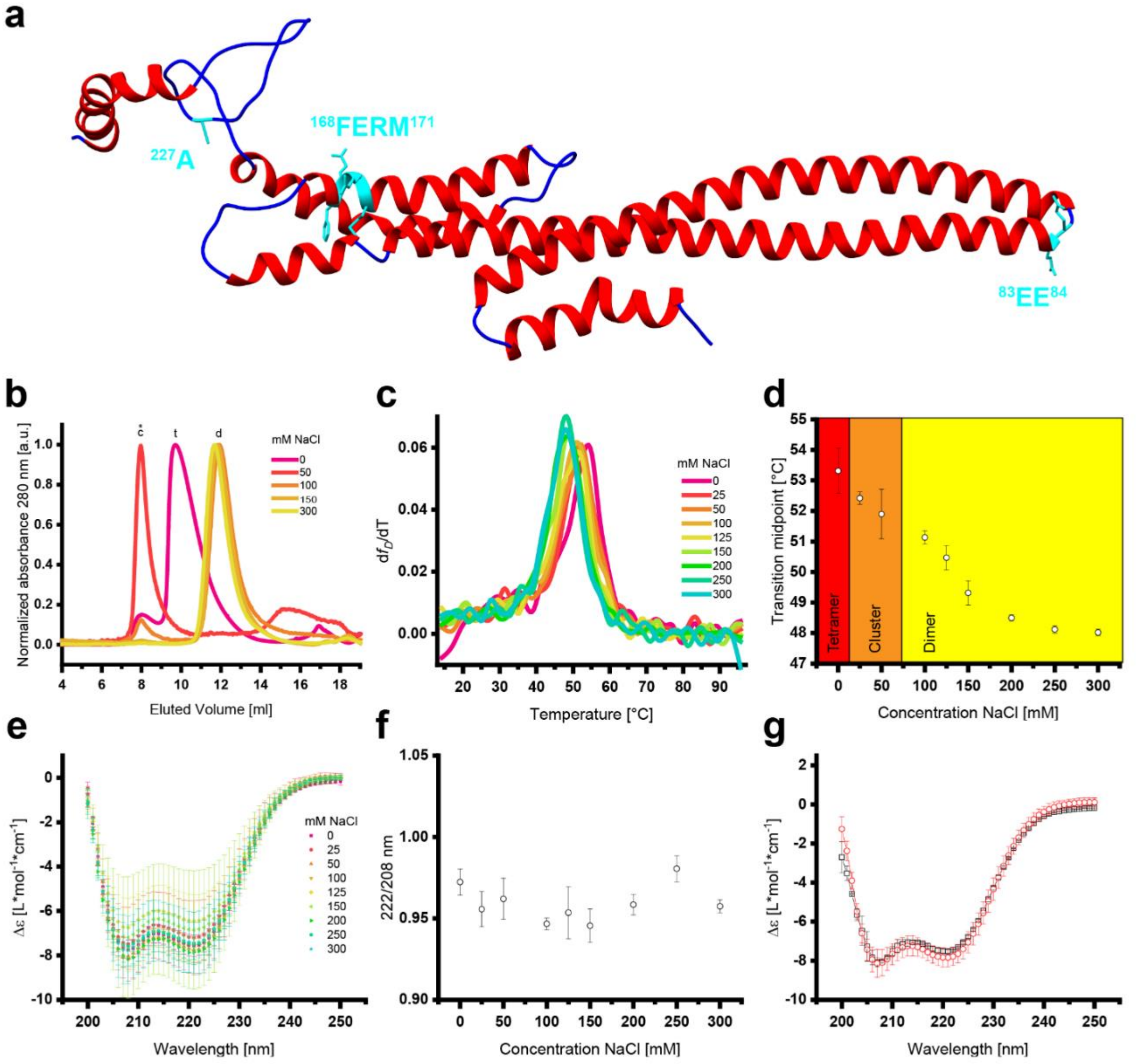


***Supplementary Figure 2:*** **The oligomeric state of IM30* is controlled by the ionic strength.**

**(a)** Predicted structure of IM30 monomers according to Saur *et al.*[3]. The "FERM", "EE" and the A227C mutation sites are marked in cyan. (Images of the structure models were prepared with CHIMERA[15].) **(b)** The oligomeric state of IM30* was monitored via SEC at increasing NaCl concentrations. An asterisk marks the void volume of the column. The elution volume of larger IM30 aggregates is marked with *c*, the elution volume of tetramers (120 kDa) is marked with *t*, and the elution volume of dimers (60 kDa) is marked with *d*. **(c)** Thermal denaturation of IM30* was monitored via CD spectroscopy at increasing NaCl concentrations. The signal at 222 nm was used as a proxy for the folding state. For better visualization of the transition point, the first derivative of the denaturation curve is shown. Each curve represents the average of three independent measurements. **(d)** Melting points derived from the thermal denaturation were plotted against the NaCl concentration and correlated to the oligomeric state as derived from SEC (Error bars represent SD, n=3). **(e)** CD spectra of IM30* at increasing NaCl concentrations indicate that the secondary structure shows no changes for up to 300 mM NaCl. The error bars represent SD, n=3. **(f)** The 222/208 ratio of IM30* does not change at increasing NaCl concentrations for up to 300 mM NaCl. The error bars represent SD, n=3. **(g)** CD spectra of IM30* before (native, black squares) and after heating to 70 °C followed by slow cooling (refolded, red circles) indicate complete reversibility. The error bars represent SD, n=3.

*AFM studies of IM30 on mica and solid supported lipid bilayers (SLBs)*

First, we imaged the structure of IM30 via AFM in absence of a lipid bilayer, as a reference for the unbound structure. Thus, we let IM30 adsorb on bare muscovite mica in 20 mM HEPES. Under these conditions, the protein formed punctae on the substrate with variable diameters and heights (Supplementary Fig. 4c/d). In 10 mM Tris buffer (pH 7.6) containing 150 mM KCl, we found IM30 forming well-defined ring-like structures with diameters ranging from 16 – 30 nm and apparent heights of 15 to 21 nm (Supplementary Fig. 3a). The dimensions of these rings are in excellent agreement with the dimensions of the 3D reconstruction of IM30 ring structures based on TEM analyses (24 – 32 nm diameter and 13 – 15 nm height)[3]. Hence, we concluded that the IM30 ring is more stable when it binds to the mica substrate in the above-mentioned Tris-buffer conditions. Yet, we also noticed that some of the ring-shaped particles looked distorted and exhibited additional material around/below them (Supplementary Fig. 4a). The abundance of such distorted structures increased upon longer incubation time on the substrate, and in some cases we even found particles on the substrate that resembled the punctae we observed in HEPES buffer. Possible reasons for the occurrence of such distortions may be dissociation over time or manipulation of the protein particles with the AFM tip during the measurement. Indeed, we found that scanning individual particles with a smaller scan area at high resolution led to flattening and loss of the defined ring shape. Thus, we were restricted to comparably large scan areas with less resolution. Nevertheless, even samples that were incubated for longer times on mica without imaging showed the additional material around/below them in the first scan (Supplementary Fig. 3b). Consequently, IM30 rings appear to disassemble upon interaction with the negatively charged mica surface. However, in contrast to DOPG bilayers, no carpet formation was observed. To exclude that the observed disassembly and subsequent carpet formation are induced by the extreme surface charge of a pure PG bilayer, we have repeated the AFM experiments with a PC(phosphatidyl choline):PG (60:40) mixture. On these surfaces we also saw ring disassembly and carpet formation (Supplementary Fig. 5).

Nevertheless, on the mica surface IM30 appeared to be more prone to ring disassembly in HEPES buffer than in Tris buffer, as we exclusively found undefined *punctae* in HEPES. Indeed, when we analyzed IM30 in Tris and in HEPES buffer via CD-spectroscopy, the structure of IM30 differed (Supplementary Fig. 4e). In HEPES buffer, we found the typical mainly α-helical structure with a coiled-coil ratio of 1.20±0.10, whereas in Tris buffer the α-helix content was reduced and the coiled-coil ratio was increased to 1.49±0.09. This difference in the structure did also affect the stability of the IM30 secondary structure against urea denaturation. In HEPES we found a transition midpoint at ~2.67 M urea. In contrast, the IM30 stability is substantially reduced in Tris buffer, indicated by a transition midpoint at ~2.44 M urea and a much shallower transition slope (Supplementary Fig. 4f). The altered secondary structure of IM30 in Tris buffer is most likely responsible for the increased stability of the ring structure upon binding to the mica substrate. However, the resistance of the secondary structure against urea denaturation was negatively influenced by the altered structure under (close to) equilibrium conditions. Thus, we decided to conduct our (AFM) experiments in HEPES buffer. Furthermore, we have found in previous experiments that the protein is not able to fulfill most of the described *in vitro* activities in the presence of Tris. In particular, IM30 does not bind to membrane surfaces anymore. Yet, while Tris buffer is not relevant for the study of the conformation of IM30 on membrane, our results show that AFM is capable of imaging IM30 with resolutions that allow discrimination between the prototypical ring structures and deviations from that (Sup Figure 3).

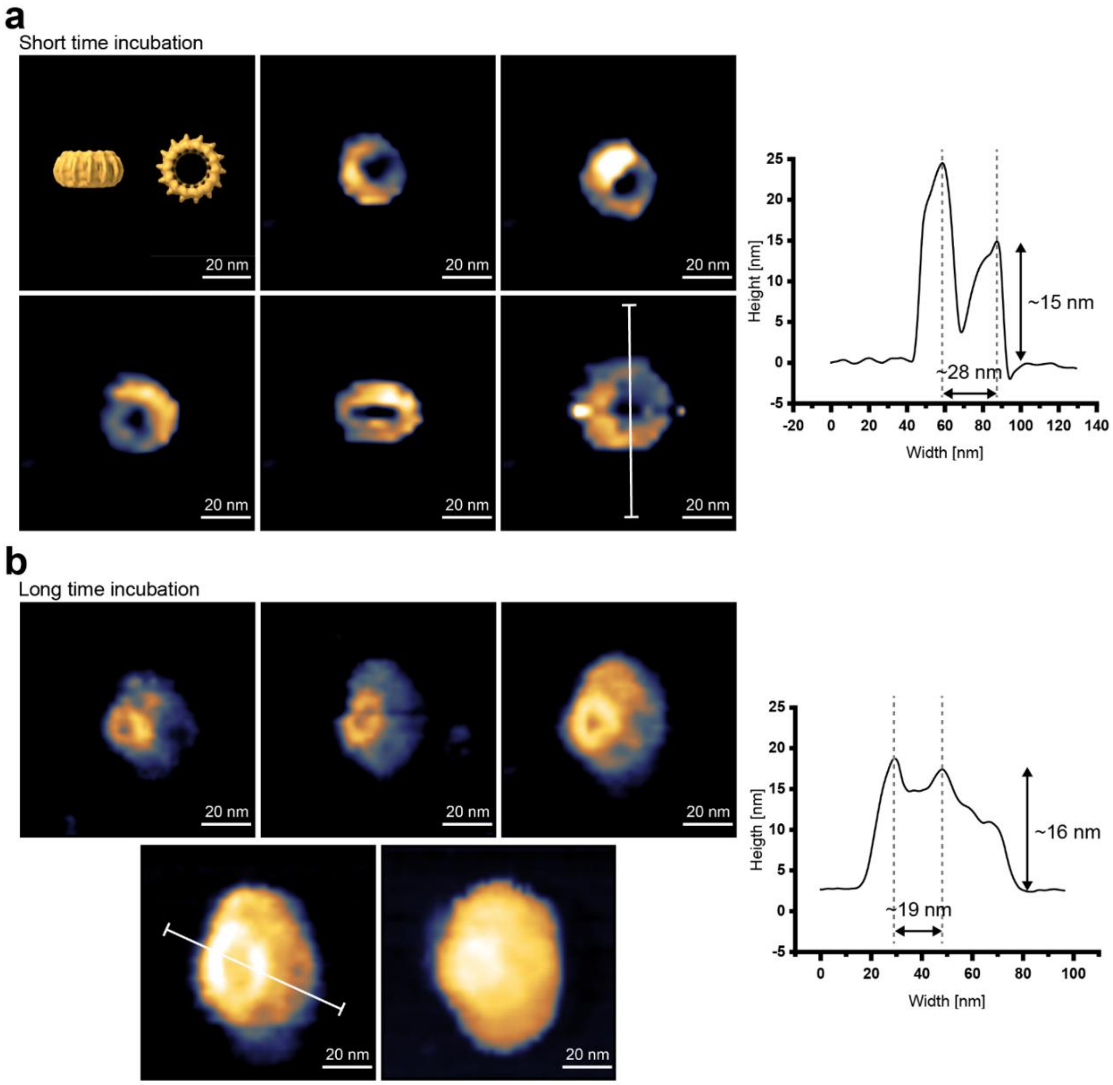


***Supplementary Figure** 3*: **IM30 binds to mica surfaces initially as a ring.**

A series of AFM height images is shown, visualizing IM30 bound to mica substrate in Tris buffer. The images were cropped to highlight individual particles. A Gaussian filter (size 1.18 px) was applied to the cropped images, and they were finally scaled to 512 px (interpolation type "Schaum"). Uncropped and unfiltered images are shown in Supplementary Fig. 4a/b. IM30 forms ring-shaped structures with variable diameters. **(a)** Upon extended incubation on the substrate, the ring-shaped structures were still detectable, but additional material around/below the rings distorted the well-defined ring-shape. **(b)** The position of the height profiles is marked by a section line. For comparison, a TEM reconstruction of an IM30 ring (EMD:3740)[3] is shown scaled to the same magnification.

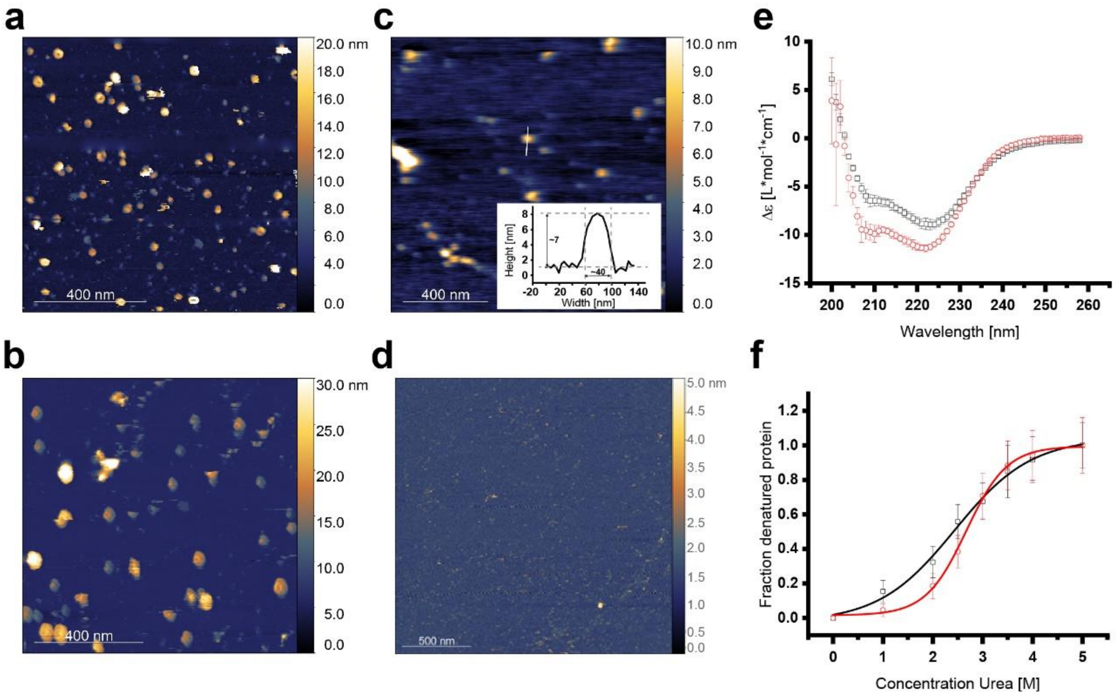


***Supplementary Figure 4*: Tris buffer affects the structure and stability of IM30.**

**(a)** The uncropped AFM height images of IM30 rings bound to mica after short incubation (~10 – 15 min) in Tris buffer are shown. Scar marks were manually masked, and the data below the mask were removed by laplacian interpolation with GWYDDION[16]. The false-color ruler shows the height scale. **(b)** The uncropped AFM height images of IM30 rings bound to mica after extended incubation time (> 1 h) in Tris buffer is shown. Scar marks were manually masked, and the data below the mask were removed by laplacian interpolation with GWYDDION[16]. The false-color ruler shows the height scale. **(c)** An AFM height image of IM30 bound to mica in HEPES buffer is shown. The section line marks the position of the height profile. The false-color ruler shows the height scale. (inlet) A height profile of the undefined IM30 particles in HEPES buffer is shown. **(d)** An AFM height image of small IM30* oligomers bound to mica in HEPES buffer is shown. No rings or carpet structures are visible. **(e)** CD-spectra of IM30 in HEPES (red circles) and Tris (black squares) buffer, respectively, are shown. Error bars represent SD with n=3 (HEPES) and n=4 (Tris). **(f)** Urea denaturation of IM30 in HEPES and Tris buffer reveals increased stability of the IM30 secondary structure in HEPES buffer. The transition midpoints ($T_m$) were determined by fitting with two-state denaturation model (Boltzmann). $T_m$ in HEPES=2.67±0.03 M ($R^2$=0.99775); $T_m$ in Tris=2.44±0.11 M ($R^2$=0.9914). Error bars represent the Gauss error propagation after normalization to folded (0) and unfolded (1) state, n=3.

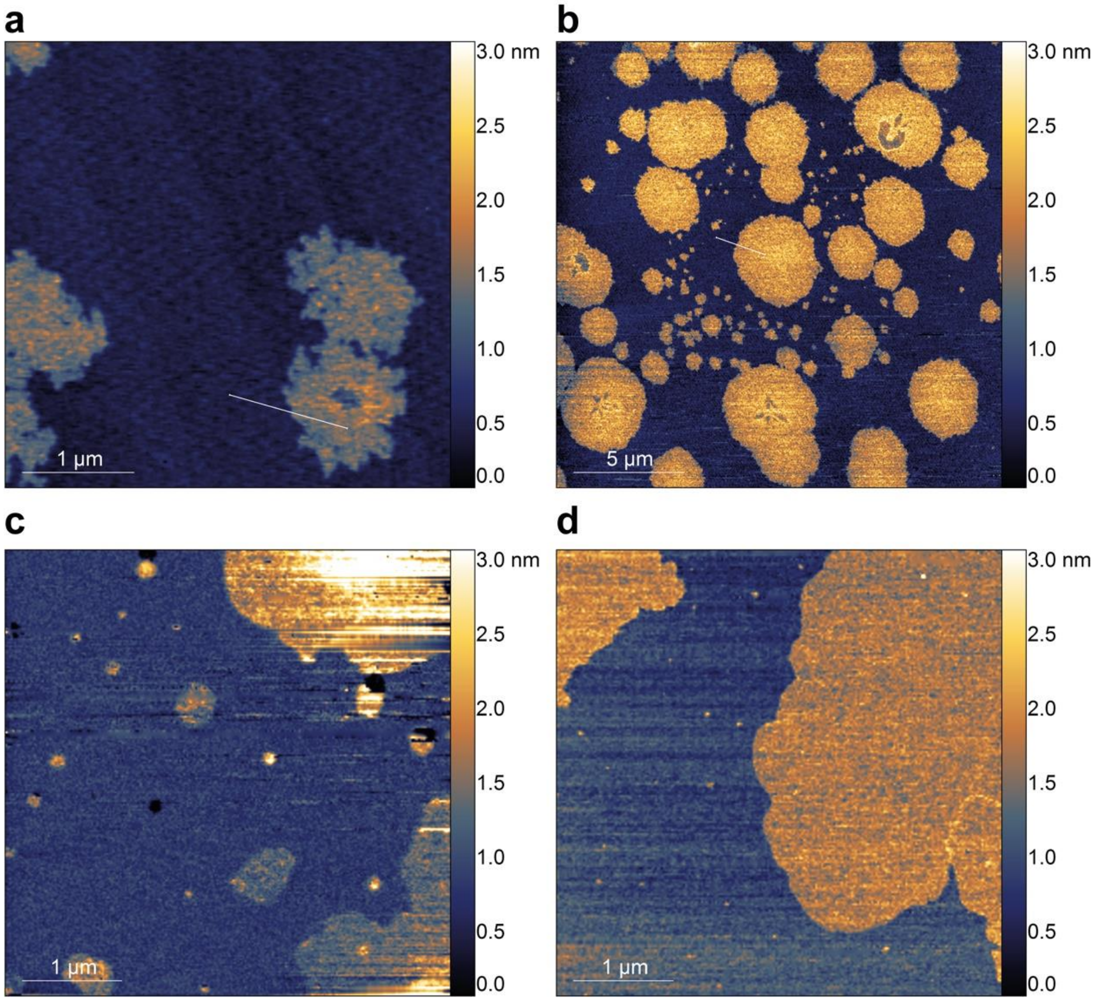


***Supplementary Figure** 5*: **AFM images of IM30 WT and IM30* on a lipid bilayer in HEPES buffer.**

**(a)** Uncropped height image of IM30 WT carpets bound to a DOPG SLB in HEPES buffer. The white line marks the region used for the height profile. **(b)** Uncropped height image of IM30* carpet structures on a DOPG SLB in HEPES buffer. The white line marks the region used for the height profile. **(c)** Height image of IM30 WT carpets bound to a SLB made of a 60:40 DOPC:DOPG lipid mixture. **(d)** Height image of IM30 * carpets bound to a SLB made of a 60:40 DOPC:DOPG lipid mixture. (The false-color ruler shows the height scale. All AFM images were prepared with GWYDDION [16]).

*Up to 10% isopropanol does not affect the structure of IM30*

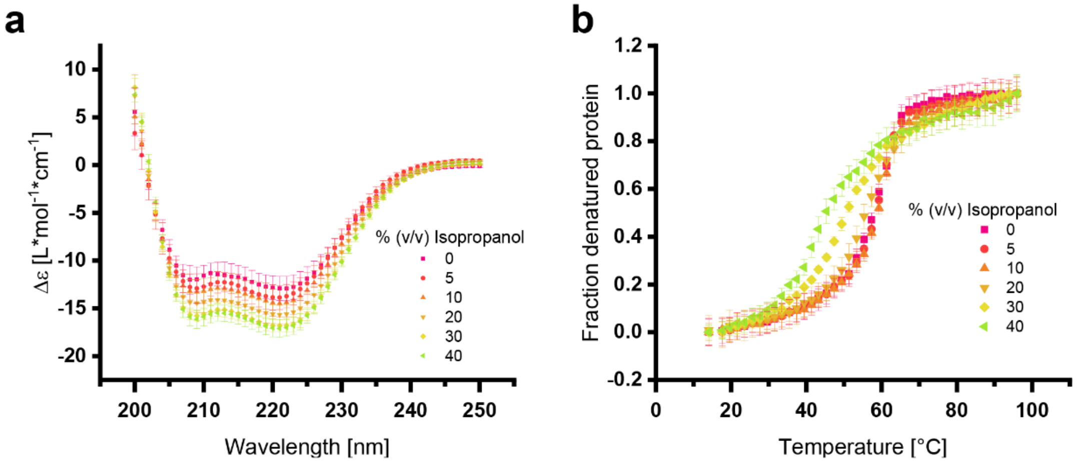


***Supplementary Figure 6*: The secondary structure and stability of IM30 does not change upon addition of up to 10% isopropanol.**

**(a)** CD spectra of IM30 WT at increasing isopropanol concentrations verify that the secondary structure shows only minor changes for isopropanol concentrations of up to 10%. The error bars represent SD, n=6. **(b)** Thermal denaturation of IM30 WT in presence of increasing isopropanol concentrations is shown. The denaturation of the protein was monitored at 222 nm. The denaturation curve does not change upon addition of up to 10% isopropanol. The error bars represent SD, n=3.

*Additional information: MS limited proteolysis*

**a**

```
Similarity : 157/267 (58.80 %)

IM30*  1     MGLFDRLGRVVRANLNDLVSKAEDPEKVLEQAVIDMQEDLVQLRQAVARTIAEEKRTEQR  60
             ############||||||||||||||||||||||||||||||||||||||||||||||||
Seq    1     ------------ANLNDLVSKAEDPEKVLEQAVIDMQEDLVQLRQAVARTIAEEKRTEQR  48

IM30*  61    LNQDTQEAKKWEDRAKLALTNGAANLAREALARKKSLTDTAAAYQTQLAQQRTMSENLRR  120
             ||||||||||||||||||||||||||||||||||||||||||||||||||||||||||||
Seq    49    LNQDTQEAKKWEDRAKLALTNGAANLAREALARKKSLTDTAAAYQTQLAQQRTMSENLRR  108

IM30*  121   NLAALEAKISEAKTKKNMLQARAKAAKANAELQQTLGGLGTSSATSAAAAAENKVLDMEA  180
             |||||||||||||||||||||||#####||||||||||||||||||||||||||######
Seq    109   NLAALEAKISEAKTKKNMLQARA-----NAELQQTLGGLGTSSATSAAAAAENK------  157

IM30*  181   TSQAAGELAGFGIENQFAQLEASSGVEDELAALKASMAGGALPGTSAATPQLEAAPVDSS  240
             ############################################################
Seq    158   ------------------------------------------------------------  157

IM30*  241   VPANNASQDDAVIDQELDDLRRRLNNL  267
             ###########################
Seq    158   ---------------------------  157
```

**b**

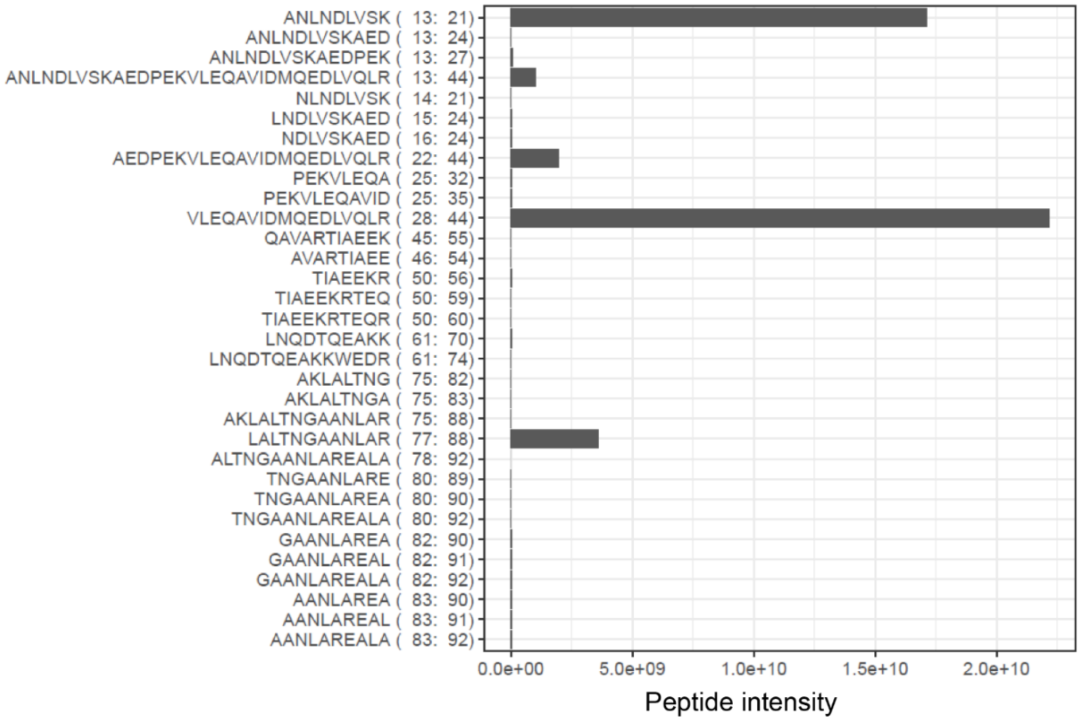


***Supplementary Figure 7*: Sequence coverage of IM30* analyzed by MS.**

**(a)** The sequence coverage of IM30* after limited proteolysis with endoproteinase GluC and analysis via in-gel digestion and MALDI/MS is shown. **(b)** The peptides identified by in-gel digestion and MALDI/MS and the respective peptide intensity are shown.

*IM30_H3b-7 is mostly unstructured*

In both, IM30 WT and IM30*, amino acids in the predicted helices 2 and 3a exhibited a dynamic behavior of deuterium incorporation that is indicative of an intact secondary structure (*i.e.*, low H/D exchange upon short incubation time of 10 s with a gradual increase over the time course of up to 10000 s (Supplementary Fig. 8a/b)), and thus in agreement with their assignment as helices 2 and 3a in the predicted model of Saur and coworkers[3]. In contrast, the predicted linker region between helices 6 and 7 exhibits high H/D exchange in both variants even after the shortest time of deuteration with no marked change over the time course, indicative of an unstructured region. The succeeding region previously assigned as helix 7 follows a similar pattern and, therefore instead appears to be mostly flexible/unstructured in both proteins (Supplementary Fig. 8a/b). In addition to the potentially unstructured regions in IM30 WT, we observed approximately 40-80% H/D exchange for the entire H3b-7 region in our HDX-MS experiments of IM30* after the shortest deuteration time (*i.e.*, 10 s) with no further increase in deuteration after prolonged incubation. This implies that the region H3b-7 is either highly flexible or unstructured (Supplementary Fig. 8a).

To test this notion, we recorded CD and 1D-$^{1}$H-NMR spectra of unlabeled IM30_H3b-7. The CD spectrum of IM30_H3b-7 shows a pronounced minimum at 202 nm which is indicative of a highly unstructured polypeptide (Supplementary Fig. 9a) in agreement with the lack of dispersion in the amide/aromatic and aliphatic regions of the 1D-$^{1}$H-NMR spectrum (Supplementary Fig. 9b/c)[17]. Thus, IM30_H3b-7, as well as the C-terminal domain of IM30*, are indeed intrinsically disordered.

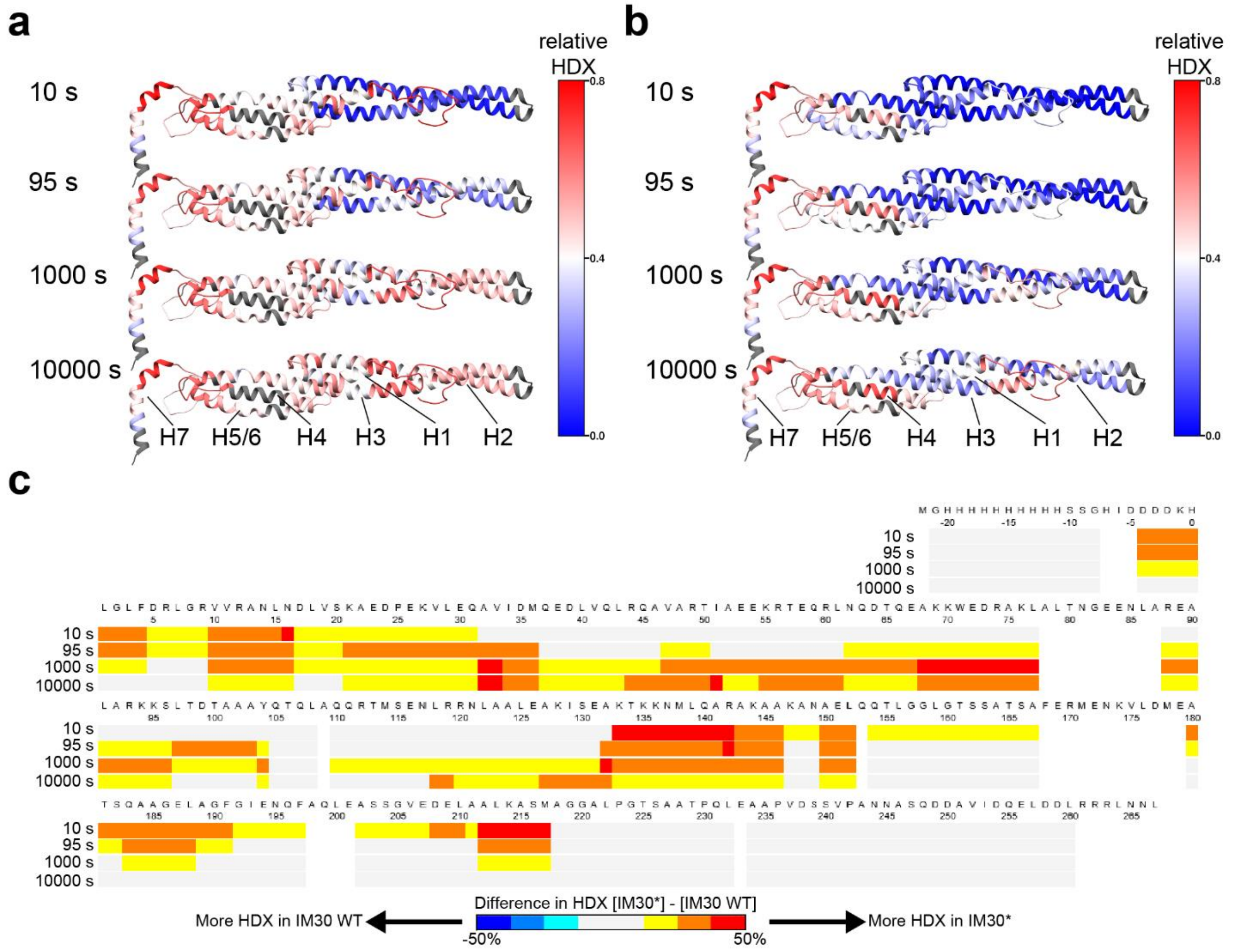


***Supplementary Figure 8*: HDX of IM30* and IM30 WT.**

**(a)** The relative H/D exchange of IM30* after incubation with a deuterated buffer for different times was mapped on the predicted IM30 monomer structure[3]. Dark grey regions mark sites where no corresponding peptides were found. **(b)** The relative HDX of IM30 WT after incubation with a deuterated buffer for different times was mapped on the predicted IM30 monomer structure[3]. Dark grey regions mark sites where no corresponding peptides were found. **(c)** The difference in deuterium incorporation between IM30 WT and IM30* is shown. (Images were prepared with CHIMERA[15] and DynamX 3.0 (Waters).)

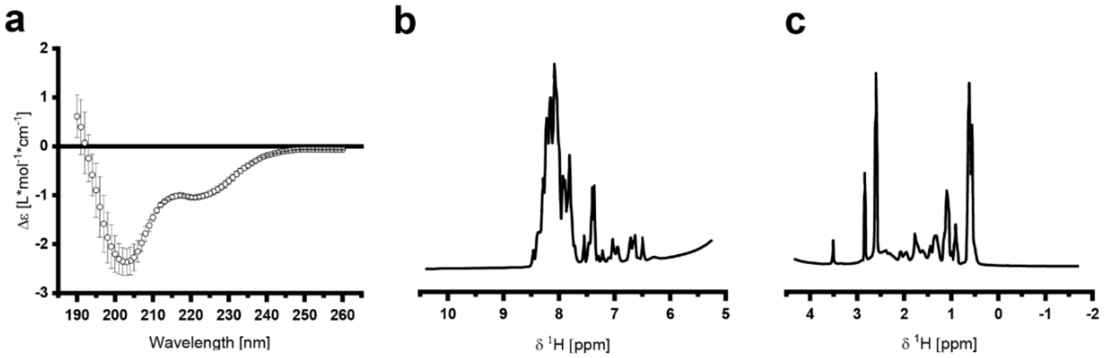


***Supplementary Figure 9:*** **The IM30* C-terminal domain is intrinsically disordered.**

**(a)** The CD-spectrum of IM30_H3b-7 reveals that the protein is mostly unstructured. Error bars represent standard deviation (n=3). (**b/c)** 1D-$^{1}$H-NMR spectrum of unlabeled IM30_H3b-7 is indicative of an unstructured protein (**b**: amide/aromatic proton region; **c:** aliphatic proton region).

*Ab initio bead model generation based on the SAXS data*

As the front region of the SEC peak in our SEC-SAXS experiment yielded a homogenous protein fraction with a molecular mass of 63.2±5.2 kDa (Supplementary Fig. 10a), we proceeded to analyze the SAXS data in order to calculate an *ab initio* bead model of the structure. The Guinier analysis of the data at low angles ($q*R_g < 1.3$) gave an $R_g$ of 6.13±0.05 nm and an extrapolated scattering intensity at zero angle of $I_0$=579.9±3.5 (Supplementary Fig. 10b). The data resulting from the Guinier analysis were used to estimate the molecular mass of the scattering particles, using the *volume of correlation* approach[18] and the *SAXSMoW* approach[19], yielding 62.9 kDa and 62.8 kDa, respectively. Both values are in excellent agreement with the calculated mass of an IM30 dimer. The pair distance distribution function of IM30* dimers has an asymmetric shape with a pronounced tailing, which is indicative of a strongly elongated structure[20] (Fig. 2c). The P(r) function decreases smoothly to zero, intersecting at a $D_{max}$ of 26 nm, indicating proper scattering data. Furthermore, $R_g$ and $I_0$ derived from the pair distance distribution analysis ($I_0$=601.3±4.5 and $R_g$=6.86±0.07 nm) reasonably well agreed with the parameters derived from the Guinier analysis. A dimensionless Kratky-plot was used to compare the scattering data obtained using IM30* and other proteins (Fig. 2d). In this plot, a well-defined, compact and spherical protein will show a symmetric peak with a maximum at $q*Rg=\sqrt{3}$, as shown for comparison using the well-characterized protein Lysozyme (SASDA96). IM30 dimers clearly do not have a well-defined, compact and spherical structure. Instead, the Kratky curve of IM30* dimers shows a steep increase until it reaches a maximum at about 5 $q*Rg$ followed by a diffuse fade out without reaching zero intensity, indicative of extended and (partially) unfolded or flexible protein regions, as can be observed by comparison with an unfolded lysine riboswitch protein (BIOISIS ID:2LYSRR) or the Plakin domain of Human plectin (SASDBC4), as an example for an extended protein shape. The Kratky curve of IM30 dimers lies in between the curves of an unfolded and an extended protein (Fig. 2d), implying an extended and somewhat flexible dimer structure, in line with the $R_g$ and $D_{max}$ values.

The $D_{max}$ and $R_g$ from the pair distance distribution (Fig. 2c) were used as starting parameters for *ab initio* dummy residue modeling with GASBOR[21]. As the program is in principle capable of fitting data up to a resolution of 5 Å or a momentum transfer of $q=4\pi\sin(\theta)/\lambda=12\ nm^{-1}$, respectively[21], the complete scattering pattern was used for fitting. We assumed a p2 symmetry of the dimer for the modeling and a number of 290 amino acids per asymmetric unit. A few initially generated models were not unambiguous, as expected when using a single scattering pattern for modeling with GASBOR. Yet, the shape of the models differed drastically. Hence, we decided to generate a statistically significant number of models to find trends in the shapes, which resulted in 115 independently generated models with acceptable correlation to the experimental scattering pattern (Supplementary Fig. 10c). These models were clustered, and the volume of all models in each cluster was averaged using DAMCLUST[22,23]. DAMCLUST was set to use p2 symmetry and to only consider backbone atoms for all bead models. In total, this approach finally resulted in 14 unique clusters, including one cluster with only two models, which was excluded from further modeling. In Supplementary Fig. 10d, each cluster is represented by its averaged volume, and the corresponding most representative GASBOR bead model is shown next to it. The overall appearance of the clusters is either W/V or S-shaped. In most clusters, the termini are long, unbranched and straight, whereas the inner parts of the models seem to differ quite strongly.

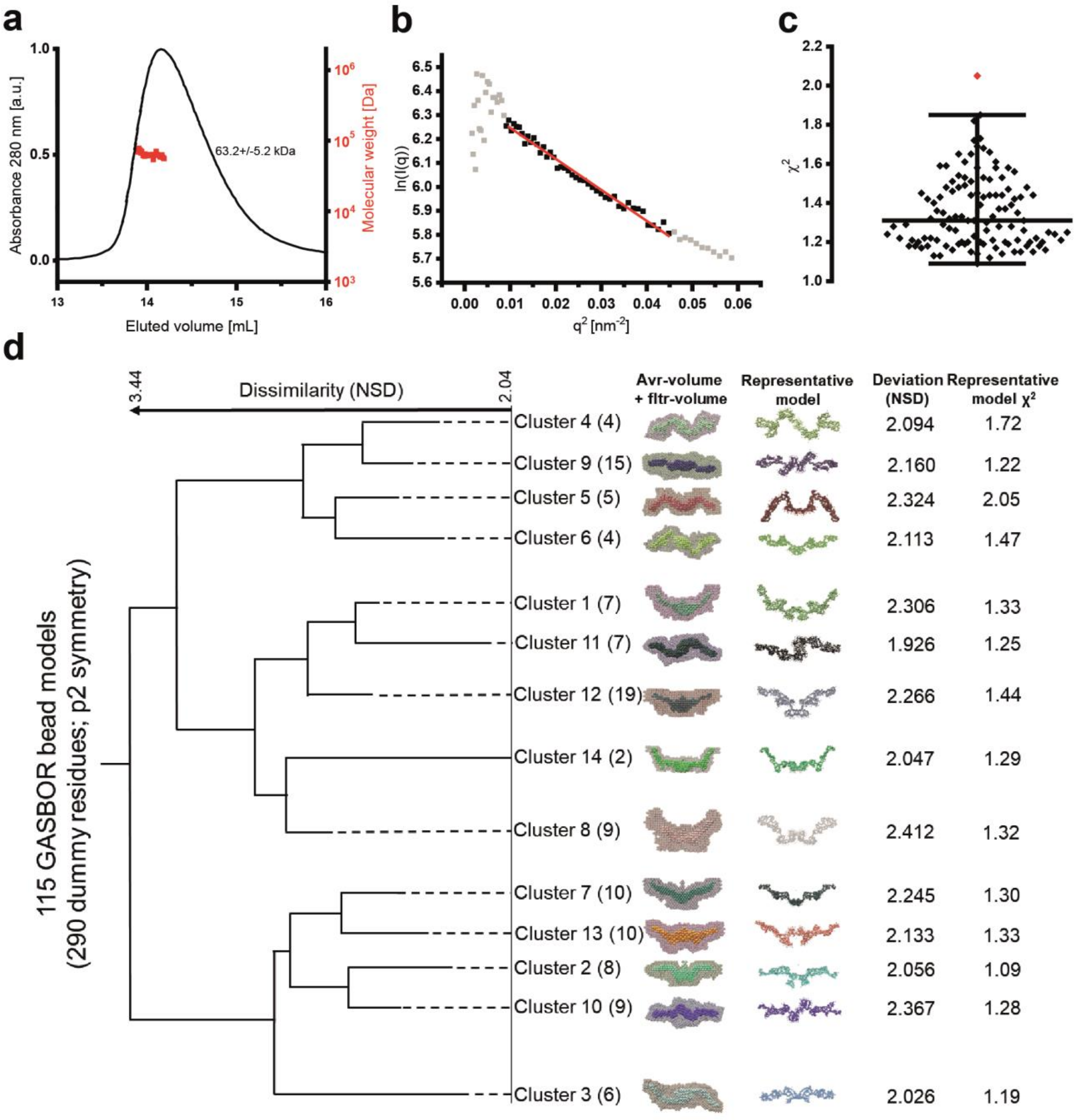


***Supplementary Figure** 10: **Ab initio*** **modeling with GASBOR resulted in 115 bead models classified into 14 clusters.**

**(a)** Only the first part of the peak was used for further analyses (indicated by red dots). The SAXS data in this range were used to estimate the molecular mass of the protein (red dots), which yields 63.2±5.2 kDa on average. **(b)** Guinier analysis of the data with $qR_g$ 0.57-1.30 gave $I_0$=579.9±3.5 and $R_g$=6.13±0.05 nm ($R^2$=0.9794). Due to their unreliability, the first 19 data points were excluded from the Guinier analysis as well as from further data processing. Except for the excluded data, no severe aggregation or inter-particle interference effects were detectable in the Guinier region. **(c)** The statistical distribution of the $\chi^2$ values of the GASBOR bead models used for the cluster analysis (n=115). **(d)** GASBOR was used to create 115 bead models that were clustered by DAMCLUST. The result of the hierarchical clustering is depicted in a similarity tree. Each cluster consist of (N) models. The average volume of each cluster (avr-volume) is shown superimposed with the filtered volume of the cluster (fltr-volume). The next columns show the most representative dummy residue bead model of each cluster, as well as the normalized spatial discrepancy (NSD) as a measure for the deviation of the representative model from the cluster. In the last column, the $\chi^2$ value of the GASBOR fit to the experimental data for each model is shown. Images of the structure models were prepared with CHIMERA[15].

*A fragmentation-based approach was used to model a structure of IM30 into the SAXS envelopes*

We aimed to build a structural model of IM30 dimers based on our SAXS envelopes. However, as only little was known about the structure of IM30 monomers/small oligomers, we again started from the predicted model described in Saur *et al*.[3]. This prediction is based on sequence-homology modeling (using the Robetta server), including the available X-ray structure of a PspA fragment as a template for helices 2 and 3 [24], rendering this part of the prediction the most reliable. Accordingly, we decided to fragment the predicted model, keeping helix 2 and 3a intact in one fragment (Supplementary Fig. 11a). This way, we also ensured that the structural core (as identified above) remains unaltered during the modeling procedure. The remaining part of the predicted model was split into five individual fragments, one for each helix. Using these fragments, we applied our modeling approach to all clusters obtained from the SAXS analysis (compare Supplementary Fig. 10d above), as described in detail in Material and Methods and Supplementary Fig. 11b.

In short, the helix fragments were fitted into the SAXS envelopes (split at the symmetry axis to obtain quasi-monomeric envelopes). To account for the unstructured regions in IM30* that were identified in the HDX experiments, we set a threshold of 45% relative HDX (after 10 s) as the limit to define a part of the structure as flexible. The flexible parts of the helix fragments were removed and MODELLER was used to recreate the missing connections between the fragments as unstructured random coils[25]. The models were refined by a simulated annealing molecular dynamics approach to fit them in the density maps created from the SAXS-cluster envelopes (FLEX-EM)[26,27], and the dimeric structure was restored by fitting two monomer models into the dimeric envelopes (Supplementary Fig. 10d). The $D_{max}$ of all models (21 to 26 nm) was in good agreement with experimentally determined $D_{max}$ (26 nm).

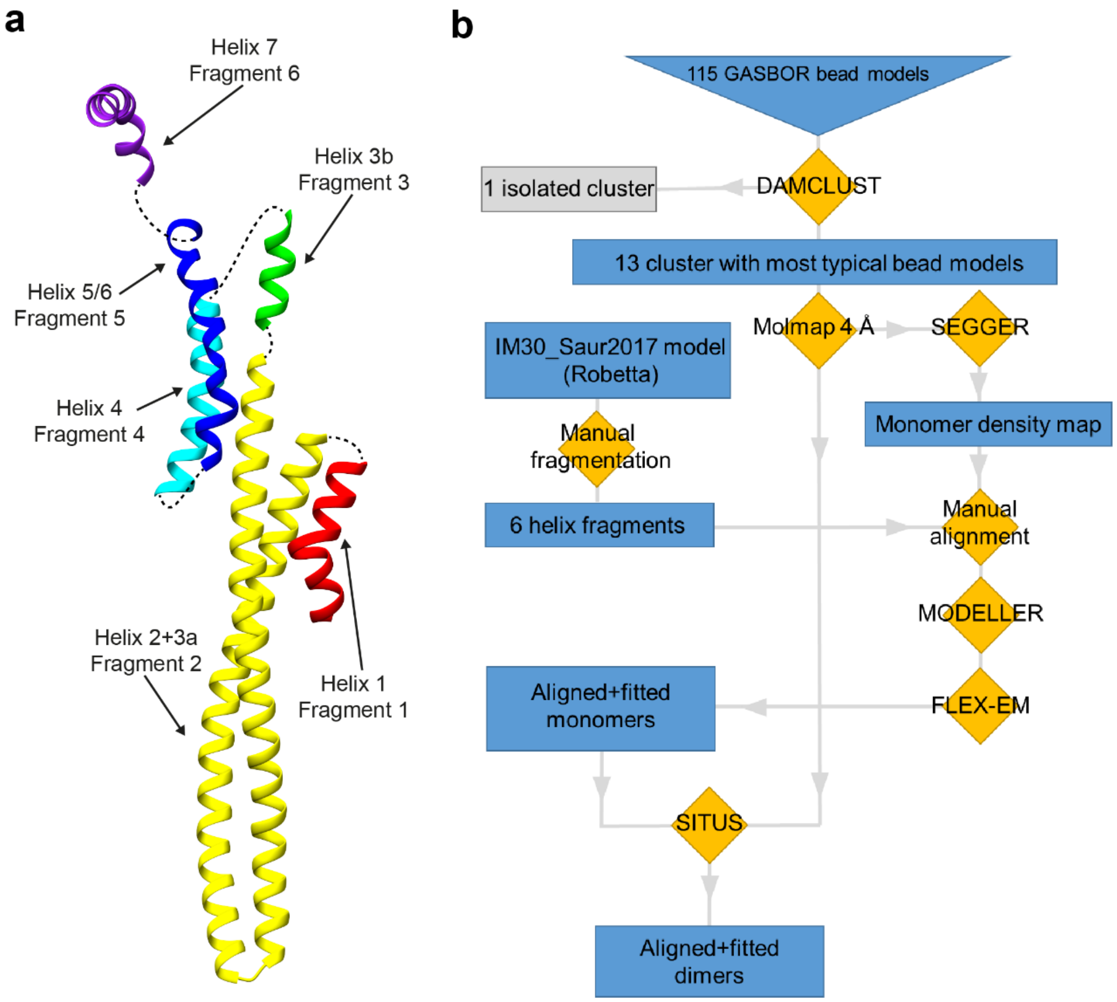


***Supplementary Figure 11:*** **Building an IM30 monomer model.**

**(a)** A predicted IM30 structure[3] was split into 6 fragments by keeping all helices intact. Images of the structure models were prepared with CHIMERA[15]. **(b)** IM30 dimers were modeled to the SAXS bead models based on a predicted structure of IM30 according to the presented scheme.

In almost all models, the structural core (helix 2/3) fitted nicely into the straight extensions of the SAXS envelopes (Supplementary Fig. 12). Thus, the dimer interface is formed by the highly unstructured C-terminal domain, which is also reflected by the high variability of the SAXS envelopes in this area. In some cases, the distance between the individual chains of the monomers in the dimer interface is relatively large, which appears quite unlikely (clusters 1, 2, and 7). Yet, in most cases, the structural overlap in the interface appears to be sufficient to provide enough surface contacts to build a stable dimer interface. However, in all cases, the flexible linker between helix 6 and 7 is mediating contact between adjacent monomers, which suggests that even in the cases with a gap in the interface the flexibility of the linker might allow close contact between the monomers.

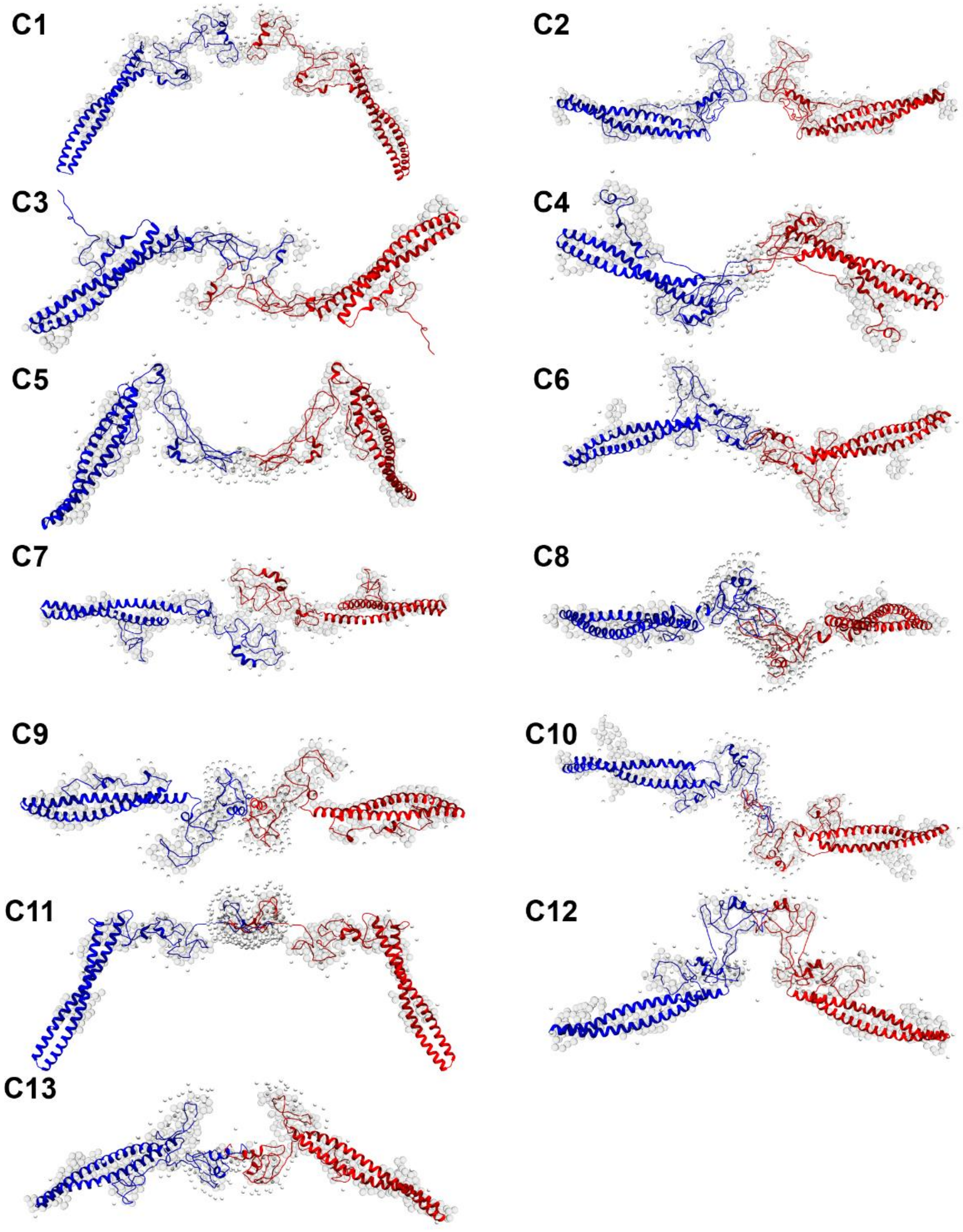


***Supplementary Figure** 12:* **A predicted IM30 structure fitted into the GASBOR bead models.**

The helical fragments were used as initial templates and fitted manually into the bead models. Afterward, the unstructured regions were replaced by random coil structures and the remaining fragments were connected by adding the missing loop structures with MODELLER. Then the structures were refined by a molecular dynamics approach using FLEX-EM. (Images were prepared with CHIMERA[15].)

*Identification of the IM30* dimer interface*

SEC coupled multi-angle laser light scattering (SEC-MALS) was used to estimate the molecular mass and the oligomeric state of two truncated versions of IM30 at buffer conditions where IM30* exclusively forms dimers. An N-terminal truncation led to IM30_H3b-7, lacking helices 1, 2 and 3a. The calculated mass of this construct is 15.2 kDa. The chromatogram of the SEC-MALS analysis is shown in Supplementary Fig. 13a. As IM30_H3b-7 does not contain any Trp residue, UV absorbance was monitored at 230 nm. The UV-absorbance signal showed only one peak at 14.4 mL, which overlapped with the second peak of the scattering signal. An additional peak in the scattering signal was found in the void volume of the column. Molecular masses were calculated using the intensity of the scattering signal and the particle concentration obtained from the UV signal. The void volume peak corresponds to masses > 1 MDa and was probably caused by minor amounts of unspecific aggregates. The second peak corresponds to a molecular mass of ~23 kDa, *i.e.* an oligomeric state of 1.51. In the case of IM30_H2-3a (Supplementary Fig. 13b), which was C-terminally truncated and lacked helices 3b, 4, 5/6, and 7 besides helix 1, we found only one major peak in both, the UV signal and the scattering signal. This peak corresponds to a molecular mass of 14.5 kDa. With a calculated mass of 16.7 kDa, we obtain an oligomeric state of 0.77 for this fragment. We conclude that IM30_H3b-7 forms dimers and IM30_H2-3a forms monomers in buffer containing 125 mM NaCl. Interestingly, SEC analyses revealed that IM30_H3b-7 forms higher oligomers (>90 kDa) when no NaCl was added to the buffer (Supplementary Fig. 13c), indicating that this domain triggers self-assembly of multiple IM30 monomers, even though it is unstructured. Taken together, we assumed that IM30* dimerizes via interactions of the C-termini.

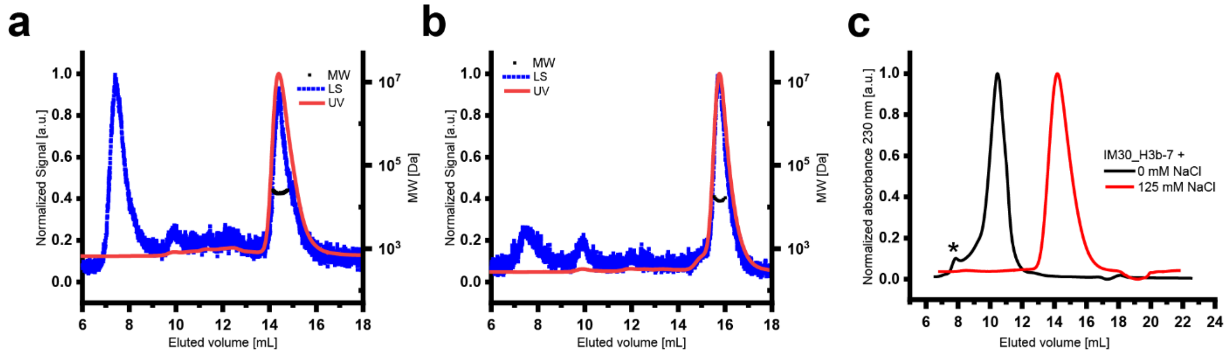


***Supplementary Figure** 13*: **The IM30* dimer interface is located in the flexible C-terminus.**

**(a):** The Multi-angle laser light scattering (MALS) analysis of a truncated IM30 construct containing only helices 3b-7 (IM30_H3b-7) is shown. The normalized scattering signal (blue dots) and the normalized absorbance at 230 nm (red line) are plotted against the eluted volume. Black dots indicate the molecular mass of the peaks, calculated from the concentration and the scattering signal. **(b):** The MALS analysis of a truncated IM30 construct containing only helix 2 and 3 (IM30_H2-3a). Data are plotted as described in A. Absorbance was measured at 280 nm. **(c):** SEC analysis of the IM30_H3b-7 construct in the absence (black) and presence (red) of 125 mM NaCl reveals that the unstructured domain (H3b-7) forms large oligomers (appararent MW >90 kDa) in the absence of NaCl and dimers in the presence of 125 mM NaCl, suggesting that the unstructured domain harbors the oligomerization interface. (The void-volume of the column is marked by an asterisk.)
